# A chromosome-scale, haplotype-resolved genome assembly of the barley stripe rust pathogen *Puccinia striiformis* f. sp. *hordei*

**DOI:** 10.64898/2026.01.25.701309

**Authors:** Rita Tam, Shideh Mojerlou, Mareike Möller, John P. Rathjen, Benjamin Schwessinger, Julian Rodriguez-Algaba

**Affiliations:** Research School of Biology, Australian National University, Canberra, Australia; Department of Agroecology, Faculty of Sciences and Technology, Aarhus University, Slagelse, Denmark

## Abstract

*Puccinia striiformis* f. sp. *hordei* (*Psh*) causes barley stripe rust, an economically important disease affecting barley production across multiple temperate regions. Unlike its well-studied wheat-infecting relative *P. striiformis* f. sp. *tritici*, genomic resources for *Psh* remain limited, with no fully documented chromosome-scale, haplotype-resolved reference genomes publicly available. Here we present a high-quality, haplotype-resolved genome assembly of the dikaryotic *Psh* isolate NP85002, generated using Oxford Nanopore long-read sequencing combined with Hi-C chromatin conformation capture. The assembly comprises 18 chromosomes per haplotype (151.3 Mb total) including 33 telomere-to-telomere chromosomes, telomeric repeats at 69/72 chromosome ends, and six internal gaps. The assembly shows high consensus accuracy (QV >72) and strong haplotype separation (97.54% within-haplotype Hi-C contacts). Genome completeness reached 90.1% complete BUSCOs, and a combined lift-over plus *ab initio* annotation achieved 94.4% complete BUSCOs. Repeat annotation indicates 45% repetitive content. We additionally provide a 102,058 bp mitochondrial genome assembly with 40 annotated genes. This genome resource provides a chromosome-scale framework for comparative and population genomic analyses of barley stripe rust and related cereal rust pathogens.

## Background & Summary

*Puccinia striiformis* f. sp. *hordei* (*Psh*), the causal agent of barley stripe rust, is an economically important fungal pathogen affecting barley production across multiple temperate regions, where epidemics result in substantial yield and grain quality losses^1,2^. The disease poses particular challenges in South Asia, East Africa, and parts of Central and North America, where *Psh* causes high disease pressure and resistance breakdown is recurrent^2-4^. As a key crop for food, feed, and malting industries^5^, barley requires durable rust control, making *Psh* a priority target for resistance breeding and integrated crop protection efforts^2,6^.

*Psh* belongs to the *P. striiformis* species complex, a group of obligate biotrophic rust fungi comprising host-adapted lineages specialized to different cereal species^4,7^. These fungi display a complex macrocyclic, heteroecious life cycle that requires two botanically distinct hosts, with haploid (n), dikaryotic (n+n), and diploid (2n) nuclear phases occurring across the sexual and asexual stages^8-10^. The rapid evolution of cereal rusts through sexual recombination, clonal propagation, and somatic hybridization (nuclear exchange without meiosis) coupled with their long-distance dispersal capacity, enables frequent breakdown of host resistance and poses persistent threats to global cereal production^11-13^. The epidemic phase on cereals is dominated by dikaryotic urediniospores that contain two haploid nuclei, which can differ markedly in sequence composition and gene content^14,15^. This divergence between the two haplotypes often leads to fragmented representations in genome assembly approaches.

Comparative studies between wheat stripe rust (*P. striiformis* f. sp. *tritici, Pst*) and *Psh* indicate that host adaptation within the *P. striiformis* complex involves gene gain and loss, structural variation, and changes in effector repertoires^16,17^. Many underlying genomic features of barley-adapted lineages remain poorly characterized, particularly regarding chromosome-level organization, telomere structure, and detailed haplotype-specific content^16^. Over the last decade, several high-quality genome assemblies have been generated for *Pst* including fully haplotype-resolved, chromosome-scale references based on long-read sequencing^18-20^. These resources are transforming our understanding of inter-haplotype diversity, structural variation and candidate avirulence genes in wheat stripe rust, providing a robust framework for population genomics and functional studies^21^. In contrast, fully documented chromosome-scale, haplotype-resolved genome assemblies for *Psh* remain absent.

Currently, only a single PacBio contig-level assembly representing a collapsed haplotype and lacking chromosome-scale scaffolding is available in public repositories^17^. Although recent efforts to generate additional *Psh* assemblies have begun, fully documented chromosome-scale, haplotype-resolved references are not yet publicly accessible^22^. High-quality, haplotype-resolved reference genomes for *Psh* are therefore essential for reference-based population resequencing and association studies targeting barley stripe rust resistance, and for enabling chromosome-level comparative and evolutionary analyses across cereal rust pathogens.

Here, we present a haplotype-resolved, chromosome-scale, and near telomere-to-telomere (T2T) genome assembly of a dikaryotic *Psh* isolate generated using Oxford Nanopore Technologies (ONT) long-read sequencing combined with chromatin conformation capture (Hi-C). We additionally provide a circular mitochondrial genome assembly. The assembly is complemented by repeat annotation, combining lift-over and *ab initio* gene prediction, and extensive quality-control analyses, providing a valuable genomic framework for future studies of *Psh* population biology, host adaptation, and effector diversity within the *P. striiformis* species complex.

## Methods

### Isolate selection and propagation

*Puccinia striiformis* f. sp. *hordei* isolate NP85002 was selected from historical collections at the Global Rust Reference Center (GRRC), Aarhus University (Denmark), and was originally collected from infected barley in Nepal in 1985. Its designation as *Psh* is further supported by host-range phenotyping reported in an accompanying study, where the isolate showed compatibility with barley lines but not with wheat lines, consistent with a barley-adapted infection profile^23^. This isolate provides a valuable temporal and geographic reference point for comparative genomics, preceding recent barley breeding efforts and documented pathogen population shifts. Fresh urediniospores were propagated on the susceptible barley cultivar Afzal under controlled greenhouse conditions (17±2 °C, 16 h photoperiod, 70-80% relative humidity). Seedlings at the two-leaf stage were spray-inoculated with urediniospore suspensions in Novec™ 7100 fluid (3M™), incubated in a dew chamber at 10 °C with >95% relative humidity for 24 h in darkness, and subsequently transferred to spore-proof greenhouse cabins^24,25^. Fresh urediniospores were harvested 15-20 days post-inoculation and stored at -80°C.

### DNA extraction and long-read genome sequencing

High-molecular-weight genomic DNA was extracted from 300 mg fresh urediniospores following the protocol described in^26^. Long-read sequencing was performed using Oxford Nanopore Technologies (ONT) on a PromethION platform. An ONT library was prepared using the Ligation Sequencing Kit v14 (SQK-LSK114) and sequenced on a single R10.4.1 flow cell with Q20+ chemistry. Raw signals were basecalled with Dorado (v0.7.2) using the super-accuracy model <u>dna_r10.4.1_e8.2_400bps_sup@v5.0.0</u> to generate simplex reads. Basecalled reads were evaluated with NanoPlot v1.44.1^27^ and seqkit v2.10.0^28^, yielding 985,067 reads and a total of 9.51 Gb, with a mean read length of **∼**9.7 kb, and read-length N50 of **∼**19.2 kb. These data correspond to **∼**60× coverage of the estimated 160 Mb diploid genome and were used for downstream *de novo* assembly.

### Hi-C sequencing

Fresh urediniospores (**∼**100 mg) were cross-linked in 1% formaldehyde at room temperature for 20 min with periodic mixing and quenched with 125 mM glycine, then washed twice in 1× PBS and pelleted, following the Phase Genomics Proximo Hi-C (Fungal) sample preparation guidelines. Cross-linked pellets were frozen and shipped on dry ice to Phase Genomics (Seattle, WA, USA). Hi-C libraries were prepared using the Proximo Hi-C (Fungal) Kit (KT6040, Protocol v4.0) with restriction enzymes DpnII, HinfI, MseI, and DdeI and sequenced on an Illumina NovaSeq 6000 (2 × 150 bp), generating 77.6 million read pairs (**∼**23.3 Gb), corresponding to **∼**145× physical coverage for chromatin contact mapping.

### Genome size estimation

Prior to genome assembly, basic genomic parameters were estimated from k-mer frequency analysis of high-quality ONT reads. K-mer counting was performed using Jellyfish v2.2.10^29^ with k = 21, and the resulting k-mer frequency histogram was analyzed with GenomeScope v2.0^30^. The analysis yielded an estimated haploid genome size of ∼75.7 Mb (diploid **∼**151 Mb), a heterozygosity rate of **∼**0.8%, and **∼**72% unique sequence content (Figure 1).

**Figure 1.**
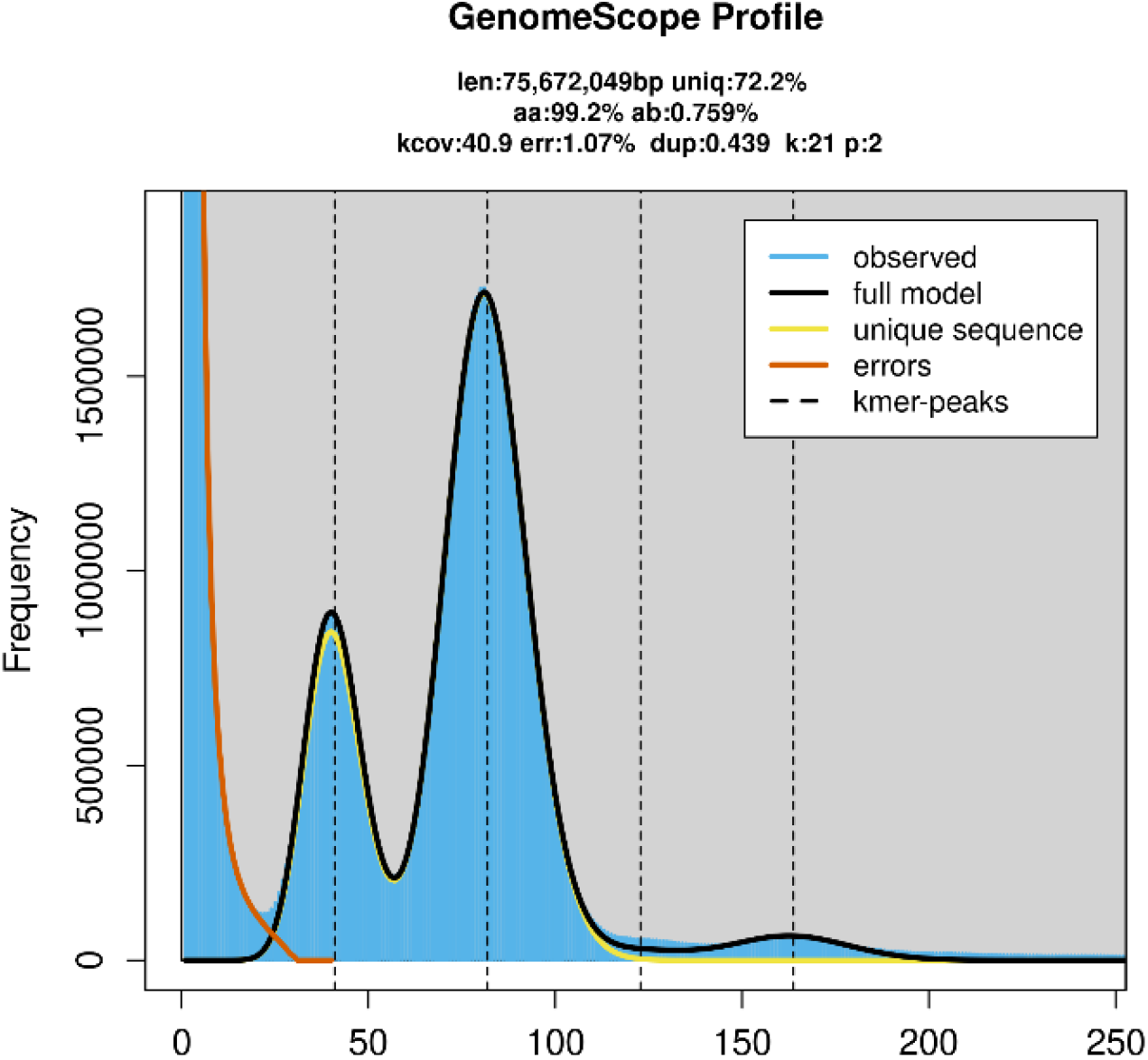
GenomeScope k-mer spectrum analysis of the barley stripe rust pathogen *Puccinia striiformis* f. sp. *hordei* isolate NP85002. The 21-mer frequency distribution and fitted model were used to estimate genome size, heterozygosity, sequencing error rate, and the proportions of unique and duplicated sequences.

### Haplotype resolved genome assembly

A haplotype-resolved genome assembly was generated from ONT and Hi-C data using hifiasm v0.25.0-r726 configured for ONT reads and Hi-C-guided phasing^31^. Simplex ONT reads were supplied with the --ont option and Hi-C read pairs with --h1/--h2. The assembler was run with the --dual-scaf to self-scaffold contigs using the homologous region on the alternative haplotype for improved contiguity, and --telo-m CCCTAA was applied to preserve telomeric repeats. This produced two haplotype-specific scaffold/contig sets (hereafter *Psh*_hap1 and *Psh*_hap2).

To remove non-nuclear and low-confidence sequences, we combined sequence-similarity and coverage filtering. All contigs were queried against the NCBI nt database using BLASTN v2.15.0+ with parameters -perc_identity 75 -evalue 1e-5 to check for contaminants and mitochondrial DNA (mtDNA). Contigs with mtDNA hits (pident ≥ 80%) were discarded. ONT reads were then mapped back to the assembly with minimap2 v2.28-r1209 (-ax map-ont -- secondary=no)^32^ and mean per-contig coverage was calculated using bamtocov v2.7.0^33^. Contigs shorter than 20 kb or with mean coverage <10× were considered low confidence and removed.

Hi-C contact maps were used to confirm nuclear haplotype phasing and to scaffold contigs into chromosomes. Hi-C reads were mapped to the cleaned assembly using Juicer v1.6^34^ and the 3D-DNA pipeline v180922^35^. Draft scaffolds were manually curated in the visualization software Juicebox Assembly Tools v2.20.00^36^. The majority of scaffolds and contigs produced by hifiasm were already at full chromosome scale where no curation was needed. The final assembly comprises 18 scaffolds per haplotype (36 in total), totaling 151.3 Mb with scaffold N50 up to 4.6 Mb, and GC content **∼**44.3% (Table 1). Chromosome naming was assigned based on homology to the Pst104E T2T genome assembly^20^. Telomere completeness was assessed by searching for TTAGGG/CCCTAA motifs within the first and last 50 bp of each scaffold using FindTelomeres.py (https://github.com/JanaSperschneider/FindTelomeres). Of the 72 expected chromosome ends (18 chromosomes × 2 haplotypes × 2 ends), 69 contained telomeric repeats; the 5ʹ ends of chr2_hap1, chr15_hap1 and chr15_hap2 lacked detectable telomeric tracts. Six assembly gaps remained in chr4_hap1/2, chr6_hap1, chr8_hap2 and chr13_hap1/2 (Table 1), most of which were associated with repetitive regions, such as the LTR retrotransposon-dense region near the *PR* locus on chr6, and ribosomal DNA repeats on chr13^20^.

**Table 1.** Assembly statistics for the *Puccinia striiformis* f. sp. *hordei* isolate NP85002 genome assembly. Metrics are reported for the combined assembly (*Psh*_hap1/2) and for each haplotype separately (*Psh*_hap1 and *Psh*_hap2).

| Assembly statistics | *Psh_hap1/2* | *Psh_hap1* | *Psh_hap2* |
| --- | --- | --- | --- |
| Total size (bp) | 151,290,331 | 75,244,898 | 76,045,433 |
| Number of T2T chromosomes | 33 | 17 | 16 |
| Scaffold N50 (bp) | 4,580,210 | 4,519,295 | 4,580,210 |
| Longest scaffold (bp) | 5,598,305 | 5,377,700 | 5,598,305 |
| GC content (%) | 44.33 | 44.32 | 44.34 |
| Telomeres detected/expected | 69/72 | 35/36 | 34/36 |
| Number of internal gaps | 6 | 3 | 3 |

### Gene prediction and annotation

Transposable elements (TEs) were predicted and annotated for each haplotype using the REPETv4 snakemake pipeline with default parameters^37^. This process identified 44.93% repetitive content in *Psh*_hap1 and 45.04% in *Psh*_hap2, dominated by Class I retrotransposons (15.78%) and Class II DNA transposons (14.97%). Identified repeats were soft-masked using bedtools maskfasta -soft^38^. Gene annotation was performed using two complementary approaches. First, a reference annotation lift-over strategy was employed. Dikaryotic evidence-based gene annotations of two high-quality *Pst* assemblies, Pst104E and AZ2, were projected onto each *Psh* haplotype using Liftoff v1.6.3^39^ chromosome by chromosome with the -chrom option^19,20^. Transferred models with invalid ORFs were filtered using agat_sp_filter_feature_by_attribute_value.pl --attribute valid_ORFs --value 0 --test “=“ (https://github.com/NBISweden/AGAT/tree/master). Second, *ab initio* prediction was run on each haplotype with funannotate predict to recover additional gene models. The Liftoff-derived and *ab initio* gene sets were merged into a single GFF3 file using agat_sp_merge_annotations.pl. Redundant CDS were deduplicated using custom Python scripts (see Code Availability). Only the isoform with the longest CDS was retained per gene, and genes encoding proteins shorter than 50 amino acids were excluded. For functional annotation, putative secreted proteins were first predicted by identifying signal peptides with SignalP v3.0 (neural network; -t euk -m nn) using the default D-score cutoff (0.433)^40,41^. To reduce false positives, mature (signal peptide-cleaved) protein sequences were screened for transmembrane domains using Phobius v1.01^42^ and TMHMM v2.0^42^. Only proteins with transmembrane predictions supported by both tools were excluded from the secretome set to ensure a conservative approach. Secondary metabolite biosynthetic gene clusters were identified with antiSMASH v8.0.4^43^. Interproscan v5.64-96.0.0^44^ was then used to assign protein domains and functional signatures, and all results were parsed with funannotate annotate to produce the final functional annotation. The final annotation comprised 17,238 protein-coding genes on *Psh*_hap1 and 17,691 on *Psh*_hap2, along with 571 and 593 tRNA genes, respectively. The mean coding sequence length was 1,076 bp with an average of 4.3 exons per gene, with a total of 4,969 single-exon protein-coding genes. These features are consistent with previously published *P. striiformis* gene structure statistics^19^. Candidate centromere regions were inferred from the Hi-C contact maps based on the characteristic “bowtie” patterns indicative of inter-chromosomal centromere-to-centromere interaction enrichment^20,45,46^. Where no clear enrichment was observed, centromere positions were estimated by aligning corresponding centromere sequences from the Pst104E isolate. Gene density was calculated in non-overlapping 10 kb windows and visualized alongside TE density and gap positions, and telomere locations using KaryploteR^47^ to provide an overview of genome organization (Figure 2).

**Figure 2.**
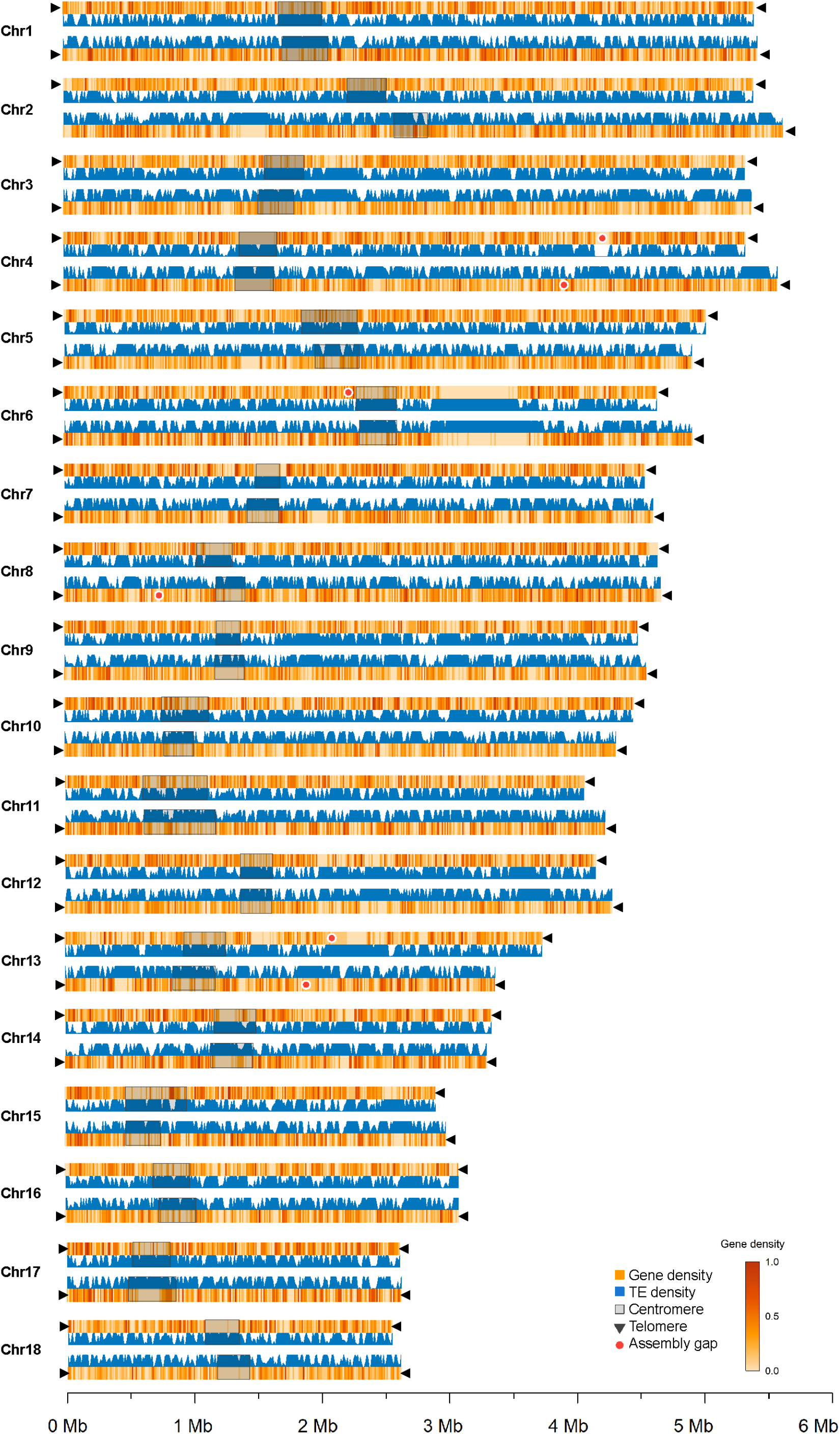
Karyotype representation of the haplotype-resolved *Puccinia striiformis* f. sp. *hordei* genome assembly. Each chromosome pair (1-18) displays haplotype 1 (upper tracks) and haplotype 2 (lower tracks). Gene density is displayed as a heatmap (orange gradient) and transposable element (TE) density is shown as a filled area plot (blue), calculated in 10 kb non-overlapping windows. Candidate centromeric regions are indicated by semi-transparent grey shaded boxes spanning both the gene density and TE tracks. Assembly gaps are marked by red circular symbols positioned on the gene density tracks. Black triangular arrowheads indicate telomeric sequences at chromosome ends, with missing telomeres absent on chromosomes 2B, 15A, and 15B. Chromosomes are ordered numerically (Chr1-Chr18) and scaled proportionally to their assembled lengths.

### Mitogenome assembly and annotation

The *Psh* mitogenome was assembled separately from the nuclear genome. Mitochondrial reads were first baited by mapping ONT reads against a concatenated FASTA file containing the *Psh* chromosome sequences and a mtDNA contig sourced from AZ2 *Pst* isolate^19^. Reads mapping to the mtDNA contig were extracted, length-filtered (20-120 kb), and assembled with hifiasm (--ont --primary), generating a circular mitogenome assembly^31^. Protein-coding and non-coding RNA genes were annotated with MFannot, an intron- and mini-exon-aware mitogenome annotator, using the genetic code table 4 (Mold, protozoan, and Coelenterate Mitochondrial; Mycoplasma/Spiroplasma)^48^. Protein sequences identified by MFannot were confirmed via BLASTP search against the ClusteredNR database. The final mitogenome assembly (total size 102,058 bp) was re-oriented to begin at the start of the *cox1* gene. A total of 40 mitochondrial genes were detected, including 14 protein-coding genes, 24 tRNA genes, one ribosomal RNA gene (*rns*) and a ribonuclease P RNA (*rnpB*) gene. The circular mitogenome assembly and annotations were visualized using the Proksee web-tool (Figure 3)^49^.

**Figure 3.**
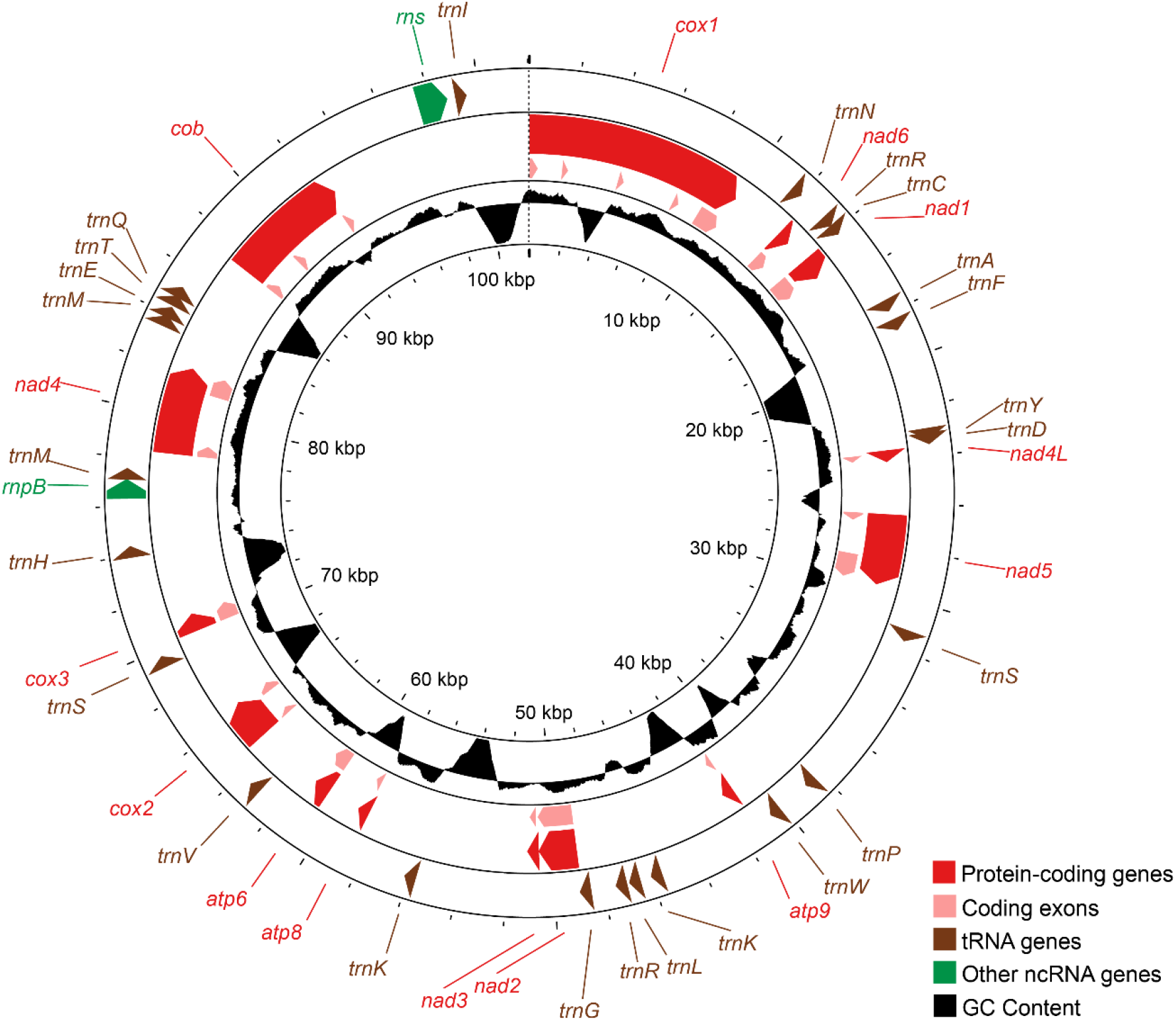
Circular representation of the mitochondrial genome assembly of *Puccinia striiformis* f. sp. *hordei* NP85002 isolate (size 102,058 bp), re-oriented to begin at the start of the *cox1* gene. The mitogenome contains 40 annotated genes, including 14 protein-coding genes, 24 tRNA genes, one ribosomal RNA gene (*rns*), and one RNase P RNA gene (*rnpB*). Protein-coding genes are shown in red, tRNA genes in brown, and other non-coding RNA genes (*rns* and *rnpB*) in green. The innermost track represents GC content variation across the mitochondrial genome.

### Data Records

All raw sequencing data generated in this study have been deposited at the National Center for Biotechnology Information (NCBI) under BioProject accession PRJNA1406799 and BioSample SAMN54803514. The Oxford Nanopore long-read and Hi-C sequencing data are available in the NCBI Sequence Read Archive (SRA) under accession group SRP665702^50^. The haplotype-resolved nuclear genome assemblies are publicly available from NCBI GenBank under assembly accessions GCA_056747125.1^51^ and GCA_056747115.1^52^, and the mitochondrial genome assembly is available under accession PX929009^53^. The genome assemblies and associated resources, including gene and transposable element annotations, are available via Zenodo (DOI: 10.5281/zenodo.18323365)^54^.

### Technical Validation

We evaluated the *Psh* assembly using k-mer-based consensus accuracy, read-mapping support, Hi-C contact structure and gene-space completeness. These complementary metrics provide an integrated assessment of assembly quality and haplotype resolution of the dikaryotic *Psh* genome.

### K-mer-based consensus accuracy

Reference-free consensus accuracy and completeness were estimated with Merqury v1.3 using a k-mer database (k = 21) built from ONT reads^55^. The combined assembly achieved a QV of 72.01 (6.30×10^−8^ error rate), with similarly high-quality values for *Psh*_hap1 and *Psh*_hap2 (QV 71.41 and 72.69, respectively). K-mer completeness exceeded 99.999% for both haplotypes with error rates of approximately 1 in 100 million bases. Spectra-cn plots showed the expected dikaryotic pattern, with most k-mers present once per haplotype and only a small fraction of higher-copy k-mers, consistent with limited haplotype collapse (Figure 4).

**Figure 4.**
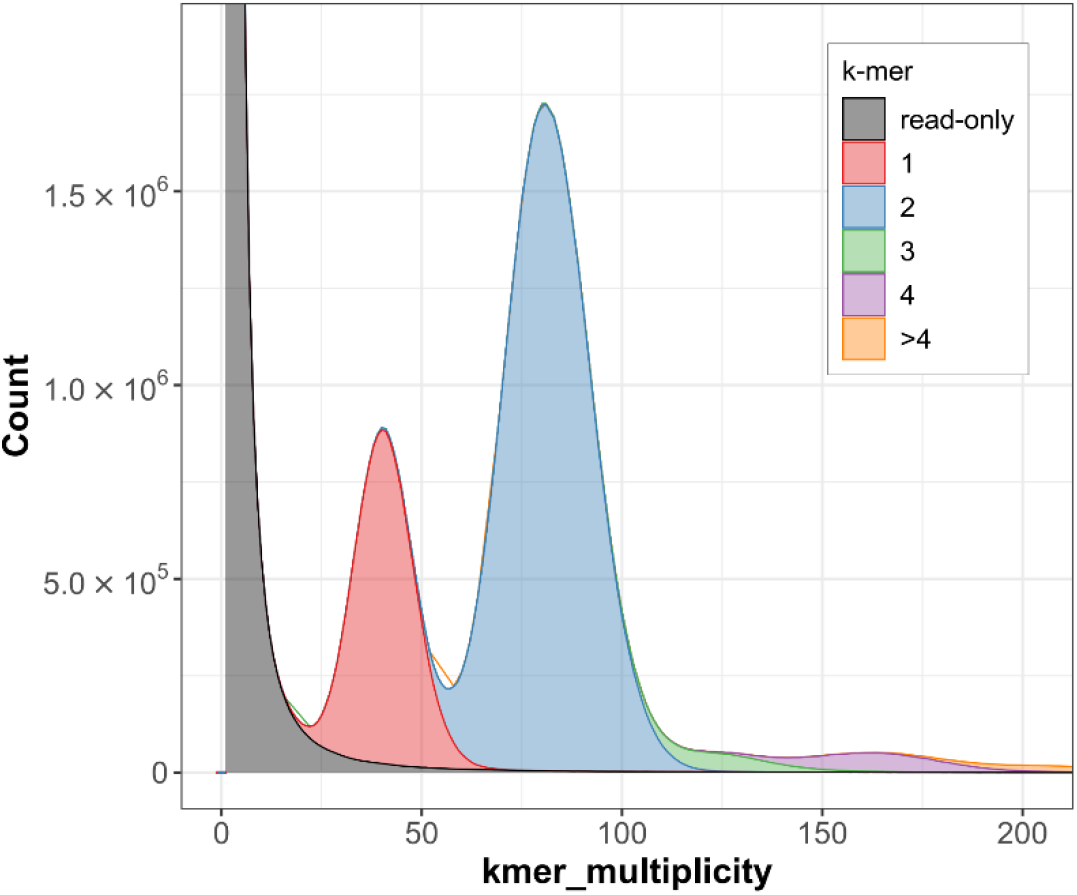
Spectra-cn plot of 21-mer multiplicity for the haplotype-resolved *Puccinia striiformis* f. sp. *hordei* assembly. The distribution shows the expected dikaryotic pattern, with most k-mers assigned to the 1× and 2× copy-number classes and only a small fraction at higher copy number (>2×), consistent with limited haplotype collapse.

### Read mapping and coverage

ONT reads were mapped back to the final assembly with minimap2 v2.28-r1209^32^ (-ax map-ont), followed by sorting, indexing, and coverage estimation with samtools v1.16^56^. Approximately 90% of ONT reads mapped to the assembled nuclear genome, with sequencing coverage ranging from **∼**98% to 100% across both haplotypes. Hi-C paired-end reads were mapped with BWA-MEM v0.7.18-r1243 to the final assembly^57^. Approximately 95% of Hi-C read pairs properly mapped, with sequencing coverage of >99%, providing robust support for chromosome-scale scaffolding and contact-map-based curation.

### Hi-C scaffolding and haplotype phasing

A Hi-C contact matrix for the full dikaryotic genome was generated using the Juicer/3D-DNA pipeline and visualized in Juicebox Assembly Tools (Figure 5). The contact heatmap clearly supported the spatial separation of homologous chromosomes into two nuclear haplotypes, *Psh*_hap1 and *Psh*_hap2. Within each haplotype, the map showed strong along-diagonal contact signals for all 18 chromosomes with no apparent off-diagonal blocks indicative of major mis-joins or misplaced contigs. Candidate centromere positions were inferred from strong inter-chromosomal contact signals and were used to confirm chromosome orientations from the shorter p-arm to the q-arm.

**Figure 5.**
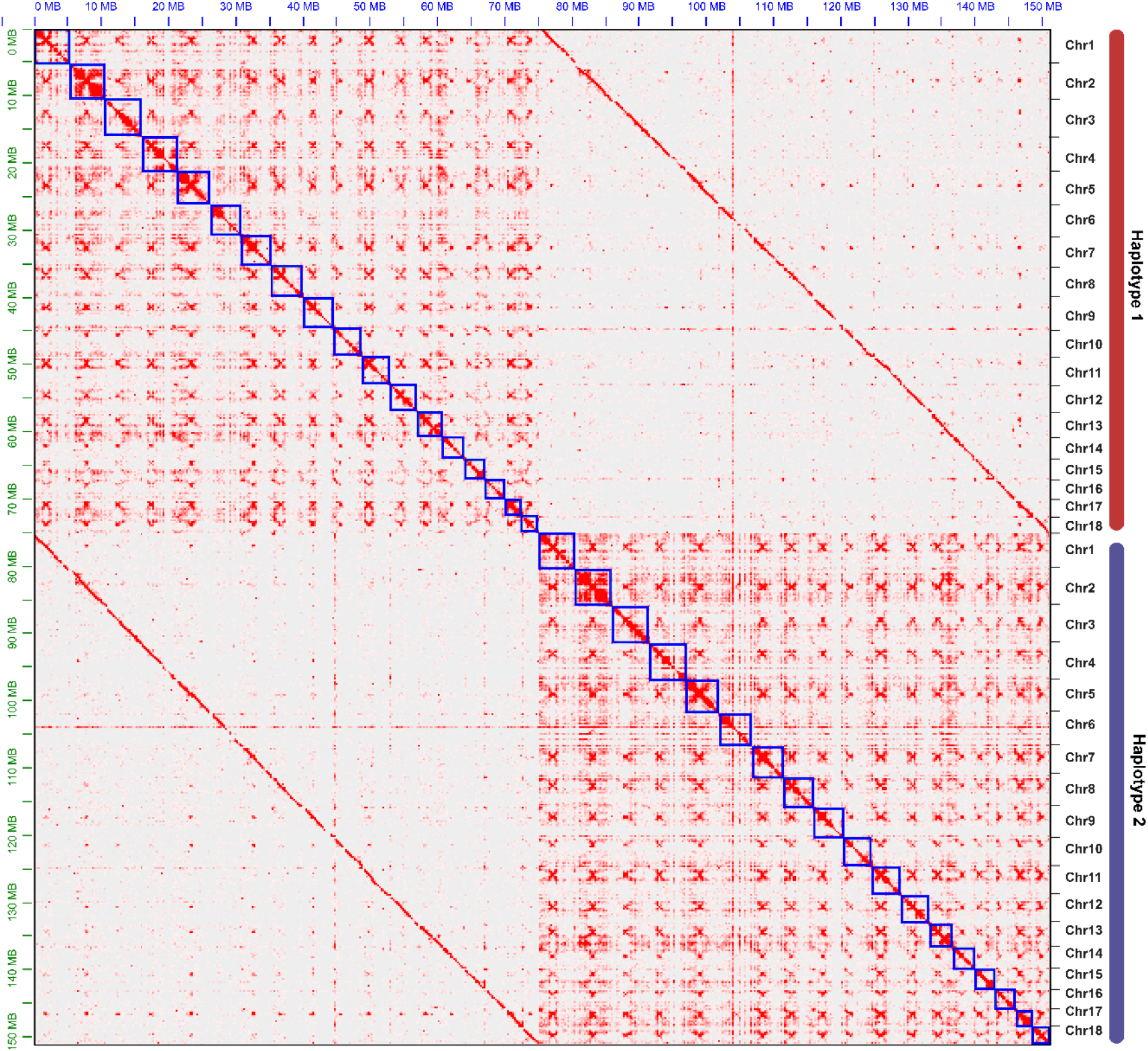
Hi-C contact map of the haplotype-resolved *Puccinia striiformis* f. sp. *hordei* assembly visualized in Juicebox. Strong within-haplotype diagonal signal across all 18 chromosomes and minimal off-diagonal structure support correct scaffolding and separation of haplotypes 1 and 2.

### Nuclear haplotype phasing correctness

To quantitatively evaluate haplotype separation and phasing accuracy, HiC-Pro v3.1.0 was used to generate a contact matrix from MAPQ≥20 Hi-C alignments^58^. *Cis-* and *trans-* chromosome contacts were quantified from raw matrix (bin size 20 kb) and visualized on a circos plot using https://github.com/RunpengLuo/HiC-Analysis^20^. The analysis demonstrated accurate phase separation with 97.54% of Hi-C contacts occurring within-haplotype contained in one nucleus, and only 2.46% representing cross-haplotype links (Table 2). The low cross-haplotype contact rate confirms successful separation of the two nuclear genomes and high-quality haplotype resolution (Figure 6).

**Table 2.** Hi-C contact counts used to assess haplotype phasing accuracy in the haplotype-resolved *Puccinia striiformis* f. sp. *hordei* assembly. Contacts were quantified from MAPQ≥20 Hi-C alignments and summarized as within-haplotype links (cis- and trans-chromosome contacts within *Psh*_hap1 or *Psh*_hap2) and cross-haplotype links (contacts between haplotypes). Values are shown as counts with percentages of total contacts.

| Hi-C contact map types | Contacts (%) |
| --- | --- |
| Within-haplotype links | 432,454 (97.54%) |
| *Cis*-chromosome, *Psh_hap1* | 176,512 (39.81%) |
| *Cis*-chromosome, *Psh_hap2* | 221,582 (49.98%) |
| *Trans*-chromosome, *Psh_hap1* | 15,158 (3.42%) |
| *Trans*-chromosome, *Psh_hap2* | 19,202 (4.33%) |
| Cross-haplotype links (total) | 10,928 (2.46%) |

**Figure 6.**
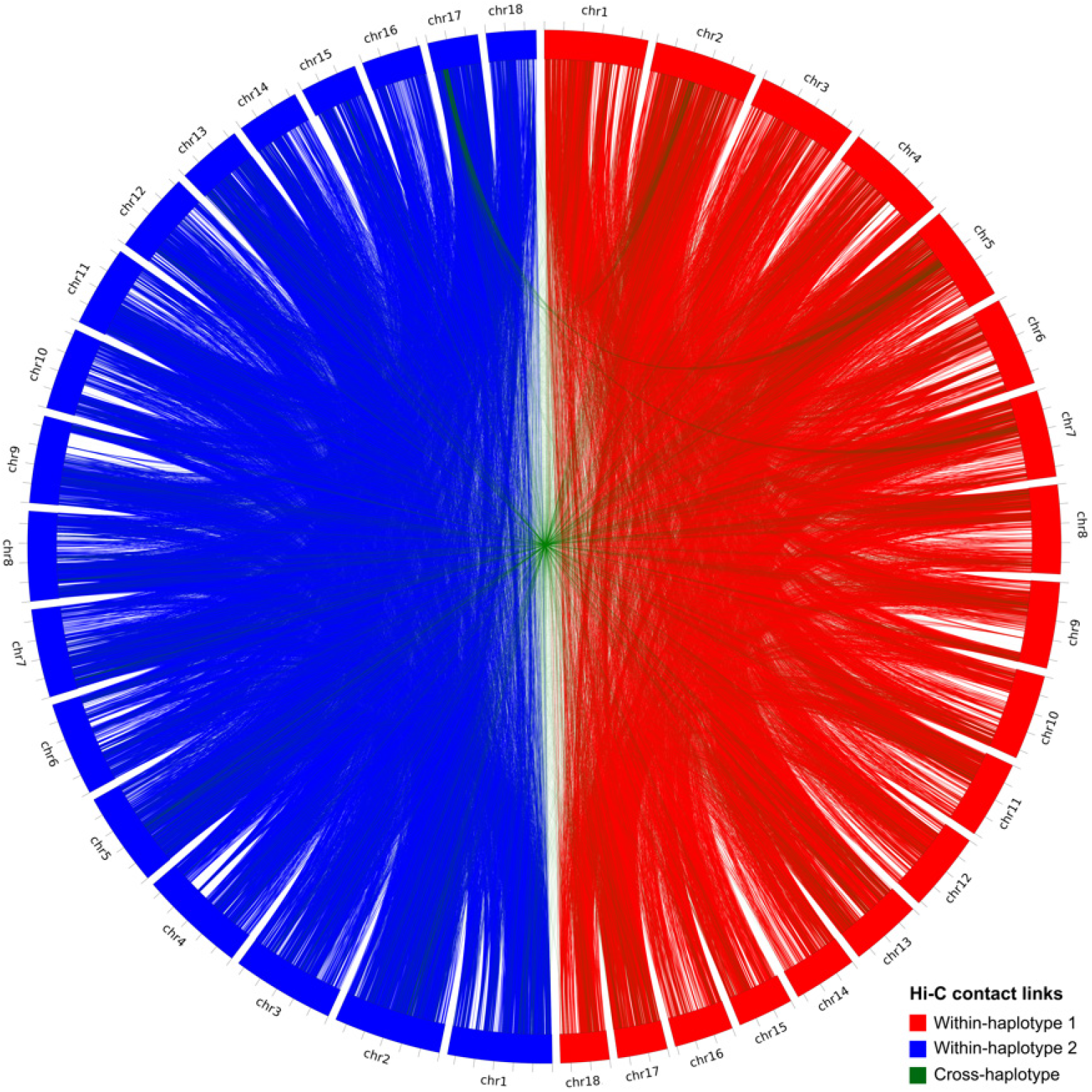
Circos plot showing within- and cross-haplotype Hi-C contact links, confirming that the two *Psh* haplotypes were accurately phased into distinct nuclei (red = haplotype 1, blue = haplotype 2). Each chromosome sequence was divided into 20 kb windows, and Hi-C contacts (MAPQ≥20) were counted per window. Cross-haplotype links between haplotype 1 and haplotype 2 are shown in green. A region on chr17_hap2 showed cross-haplotype contacts with chr2_hap1, chr5_hap1, and chr7_hap1; these contacts overlapped the estimated centromeric regions and likely reflect mapping noise associated with repetitive centromeric sequences.

### Gene-space completeness

Assembly completeness was assessed using BUSCO v6.0.0 with the basidiomycota_odb12 lineage dataset^59^. In genome mode, haplotype 1 and haplotype 2 each showed high completeness (90.9% and 91.0% complete BUSCOs, respectively), with single-copy BUSCOs comprising 89.0% and 86.9% of each haplotype, respectively. When both haplotypes were analyzed together, overall completeness was 90.1%, with most BUSCOs classified as duplicated (85.7%), consistent with the expected pattern for a haplotype-resolved assembly where orthologous genes are present in both haplotypes.

For gene annotation, lift-over projections from Pst104E (87.8% complete BUSCOs; 28,436 genes) and AZ2 (92.4%; 26,336 genes), together with *ab initio* gene predictions (86.1%; 19,084 genes), identified a higher number of BUSCOs when combined (94.4%; 34,929 genes). This supports the effectiveness of integrating lift-over and *ab initio* annotation approaches to resolve gene models despite the high repeat load and dikaryotic genome structure (Table 3).

**Table 3.** BUSCO completeness assessment of protein sequences annotated in the *Psh* genome assembly. BUSCO v6.0.0 (basidiomycota_odb12) was run on the combined assembly (*Psh*_hap1/2) and each haplotype (*Psh*_hap1 and *Psh*_hap2). Values report the percentage of complete BUSCOs (split into single-copy and duplicated), as well as fragmented and missing BUSCOs.

| Haplotype | Complete (%) | Single-copy (%) | Duplicated (%) | Fragmented (%) | Missing (%) |
| --- | --- | --- | --- | --- | --- |
| *Psh_hap1/2* | 94.4 | 2.2 | 92.2 | 1.4 | 4.2 |
| *Psh_hap1* | 93.6 | 89.0 | 4.5 | 1.5 | 5.0 |
| *Psh_hap2* | 93.5 | 86.9 | 6.6 | 1.5 | 4.9 |

## Code availability

Analyses were performed using the software and workflows described in the Methods section. Where relevant, command-line options and key parameters are reported; otherwise, tools were run using standard or recommended settings. Custom scripts used for data processing and figure generation are available on GitHub at: (https://github.com/ritatam/PshNP85002-genome-assembly).

## Acknowledgements

We thank Ellen Jørgensen and Jakob Sørensen (GRRC, Aarhus University) for assistance with isolate handling and plant multiplication, respectively. The *Psh* NP85002 isolate was part of the ‘Stubbs collection’ provided by Aarhus University, GRRC, Denmark and Plant Research International, Wageningen, the Netherlands, maintaining the Global Yellow Rust Gene Bank of the late ir. R.W. Stubbs up to 25 January 2010.

## Author contributions

R.T., B.S., and J.R.-A. designed and conceptualized the study. J.R.-A. acquired funding and supervised the research. M.M. conducted ONT long-read sequencing. S.M. extracted high-molecular-weight DNA and prepared fungal spores for Hi-C library preparation. R.T. and J.R.-A. performed bioinformatic analyses including genome assembly, annotation, quality control, and data visualization. J.P.R. and B.S. contributed to interpretation of results and provided critical feedback on the manuscript. R.T. and J.R.-A. wrote the original draft manuscript. All authors contributed to manuscript revision, reviewed the final version, and approved it for publication.

